# Binding in visual working memory is task dependent

**DOI:** 10.1101/2023.11.01.565116

**Authors:** Ruoyi Cao, Leon Y. Deouell

**Affiliations:** Edmond and Lily Safra Center for Brain Sciences, The Hebrew University of Jerusalem, Jerusalem 9190401, Israel; Department of Psychology, The Hebrew University of Jerusalem, Jerusalem 9190401, Israel

**Keywords:** Visual working memory, Binding problem, Object representation, Task dependency

## Abstract

Working memory is a neurocognitive system for maintaining and manipulating information online for a short period after the source of information disappears. The information held in working memory has been shown to flexibly match current functional goals. Considering this, we revisited the question of whether information is held in WM as separate features or as bound objects. We conjectured that rather than having a fixed answer, the format in which information is maintained in working memory is also task dependent. In two separate experiments, we investigated the binding between features when the location was not (Experiment 1, color and orientation binding) or was (Experiment 2, color and location binding) a task-relevant feature in a delayed (yes/no) recognition task by manipulating the relative relevance of conjunctions and separate features. Each experiment included two conditions, Binding dominant (BD), which emphasizes the retention of binding between features, and Feature dominant (FD), which emphasizes the retention of individual features. In both experiments, we found that memory for conjunctions was better in the BD condition and that memory for separate features was better in the FD condition. These patterns suggested that the formats of objects in VWM could be shaped by the tasks they serve. Additionally, we found that the memory of location was impaired when conjunction between location and color was task-irrelevant, while memory of color and orientation was relatively independent of each other. We conclude that the representation format of objects in working memory is influenced by task requirements.

## Introduction

Visual working memory (VWM) refers to the function of preserving and manipulating visual information for a short duration when physical stimuli are absent from direct perception (Baddeley, 1992; Esposito & Postle, 2015; Goldman-Rakic, 1995). Studies in both animals (Gilad et al., 2018; Rao, Rainer, & Miller, 1997) and humans (Myers, Stokes, & Nobre, 2017) regard VWM as a dynamic system that prioritizes task-relevant information rather than as fixed short-term storage. This was demonstrated, for example, by studies showing that VWM can selectively maintain task-relevant locations (Griffin & Nobre, 2003) or other features according to task demand (Park, Sy, Hong, & Tong, 2017; Pertzov, Bays, Joseph, & Husain, 2013; Ye, Hu, Ristaniemi, Gendron, & Liu, 2016). In this study, we investigate the flexibility of another level, namely, whether separate features or bound objects are the units of storage is task-dependent, which we refer to as the storage format below.

Studies investigating the storage format of objects in visual working memory that rely on the notion of limited VWM capacity have yielded conflicting results. Specifically, some studies have suggested that the number of objects (conjoined features), rather than the number of features, is the limiting factor (Luria & Vogel, 2011; Woodman, Vogel, & Luck, 2001a), while others have found that the number of features limits performance (An, Gong, McLoughlin, Yang, & Wang, 2014; Delvenne & Bruyer, 2004; Fougnie, Asplund, & Marois, 2010; Oberauer & Eichenberger, 2013; Olson & Jiang, 2002). One of the explored factors is whether the relevant features come from the same dimension. According to the multiple-resources view proposed by Wheeler and Treisman (2002), VWM maintains features from different feature dimensions (e.g., color and orientation) in parallel, with no additional cost, while features from the same feature dimension (e.g., two colors) compete for storage space (Wang, 2017; Wheeler & Treisman, 2002). In this study, we will focus on binding between features from different dimensions.

There is contradictory evidence regarding the retention of orientation and color in VWM. Some studies found similar memory performance for orientation and color conjunctions as for the memory of only one dimension, suggesting conjoined maintenance (Luck & Vogel, 1997). Others found that when participants were asked to recall both the color and orientation of an object, errors in memory for one feature did not significantly affect memory for the other feature, suggesting separate storage (Fougnie & Alvarez, 2011). However, while the quantity of recalled objects was largely unaffected by increasing the number of features, the precision of these representations dramatically decreased (Fougnie et al., 2010).

Spatial location information may have a special status in regard to object processing. Theoretical considerations suggest that space is extrinsic to the object, and thus, object-to-location binding is believed to mediate the binding of other features (Fougnie & Marois, 2009; Schneegans & Bays, 2017). According to feature integration theory, spatial attention is thought to be serially directed to a “master map”. At any given moment, one of the locations on the map was selected by the “window of attention”. Features located within the window are then assembled in a single object file for further analysis and identification (Treisman, 1977). Like binding between two non-location features, mixed results were obtained when binding with location was involved. Some studies have suggested that objects are automatically bound to their locations. This claim is supported by the above-chance decoding accuracy of task-irrelevant location (Foster, Bsales, Jaffe, & Awh, 2017), impaired recognition performance when memoranda appears at a location different from where it was encoded (Hollingworth, 2007; Olson & Marshuetz, 2005), and the activation of retinotopic organization in visual regions during memory delay (Beeck & Vogels, 2000; Dicarlo & Maunsell, 2003). However, other studies have provided evidence for separate representations. Lesion studies (Darling & Logie, 2006), multiple object tracking tasks (Kondo & Saiki, 2012), and behavioral interference studies (Darling, Della, & Robert, 2007) have shown a dissociation in performance between location and identity memory. Studies have also shown that memory for bound objects could be impaired without impairment in memory for individual features or locations in normal participants (Pertzov, Dong, Peich, & Husain, 2012), patients with a variant of encephalitis (Pertzov, Miller, et al., 2013) and patients with Alzheimer’s disease (Liang et al., 2016).

There are a few hints in the literature to suggest that the heterogeneity of tasks might at least partially explain some of the different results, as different tasks make conjunctions more or less relevant. Regarding the conjunction of location with objects, one study (Ester, Serences, & Awh, 2009) found that the decoding of items from BOLD signals was similar between the two hemispheres, ipsilateral and contralateral to the object held in memory, suggesting that visual details were held in VWM through spatial global recruitment of the early sensory cortex. However, by making the location task relevant, Pratte and Tong (2014) found that decoding accuracy for remembered items was greater on the contralateral side than on the ipsilateral side.

Thus, we conjectured that the storage format (separate features or conjunctions) is flexible, and it is the task that determines how information is maintained in VWM. We focus on features from separate dimensions and investigate the effect of task requirements on feature binding with (color-location) and without (color-orientation) the involvement of spatial location. In contrast to studies that manipulated the relevance of locations or other features, we were interested in directly manipulating the relevance of conjunctions.

We conducted two experiments to investigate the adaptivity of maintenance formats in WM for binding between features with and without the location as a task-relevant dimension. We surmised that even without explicitly informing subjects about the preferred mode of maintenance, the format of items in working memory will be flexibly adjusted to optimally perform the task. In a delayed (yes/no) recognition task, participants were presented with two items to be remembered. Subsequently, they had to indicate whether a probe, presented after a delay, had been shown in the initial array. Two types of probes were used, Binding probes and Feature probes; Binding probes require the memory of conjunction between features, whereas feature probes do not. The experiment contained two separate blocks with different proportions of Binding probes and Feature probes: Binding-Dominant (BD) and Feature-Dominant (FD) blocks. Consequently, conjunction memory between color and orientation was more task-relevant in the BD condition than in the FD condition. We predicted an interaction between Condition (BD and FD) and probe type (Binding probe and Feature probe), such that performance for Binding probes will be better in BD condition, while performance for Feature probe will be better in FD condition.

### Experiment 1

The aim of this experiment was to test whether task requirements could influence working memory retention of conjunctions of two non-location features, color and orientation. We tacitly manipulated the optimal strategy by presenting more frequently either probes requiring conjunction information (Binding probe) or probes requiring only feature information (Feature probes). We expected to find that when conjunctions are tested more frequently, items in working memory will be maintained as bound objects, improving conjunction recognition, whereas when separate features are tested more frequently, items in working memory will be maintained as separate features, impairing conjunction recognition.

## Materials and Methods

### Participants

Thirty-one students were recruited through the Hebrew University online study recruitment system (age range 21-36 years, mean± SD =24.79 ± 2.45, 80% females). Twenty-four of them passed the inclusion criteria described later. All participants reported no neurological ailments. The study was approved by the ethics committee of the Social Sciences Faculty at the Hebrew University of Jerusalem, Israel. Participants downloaded the computer program made with the E-prime 3 online version (https://pstnet.com/eprime-go/) and performed the experiment on their own computers. Each participant conducted the two conditions of the experiment, each on different dates within a week. They received either two course credits or a monetary compensation of 60 NISs (∼$15) for their participation.

### Stimuli

The items were Gabor gratings (5 cycles, contrast 0.7; Figure 1). The exact visual angle and size of the stimulus varied as participants performed the tests on their own screens. Each memory array comprised two Gabor gratings, with each grating having a diameter of 300 pixels. One grating was positioned on the right side of the screen, while the other was located on the left side. The distance between the center of each Gabor grating and the screen center was 200 pixels. Each grating was assigned a color from a set of four distinct colors: red (255, 0, 0), green (0, 255, 0), magenta (255, 0, 255), and blue (0, 0, 255) with an orientation out of four possible angles (3, 49, 95, or 141 degrees relative to the horizontal line). No two items within the same array had the same color or orientation, resulting in a total of 144 unique combinations. A subset of 108 of these combinations was randomly selected without repetition and used consistently across all subjects. The stimuli were created using Psychotoolbox-3, implemented in MATLAB 2018 (The MathWorks, Inc., Natick, Massachusetts). Unlike the memory array, each probe contained only one grating. There were two types of probes (Figure 1b): Binding probes and Feature probes:

**Figure 1.**
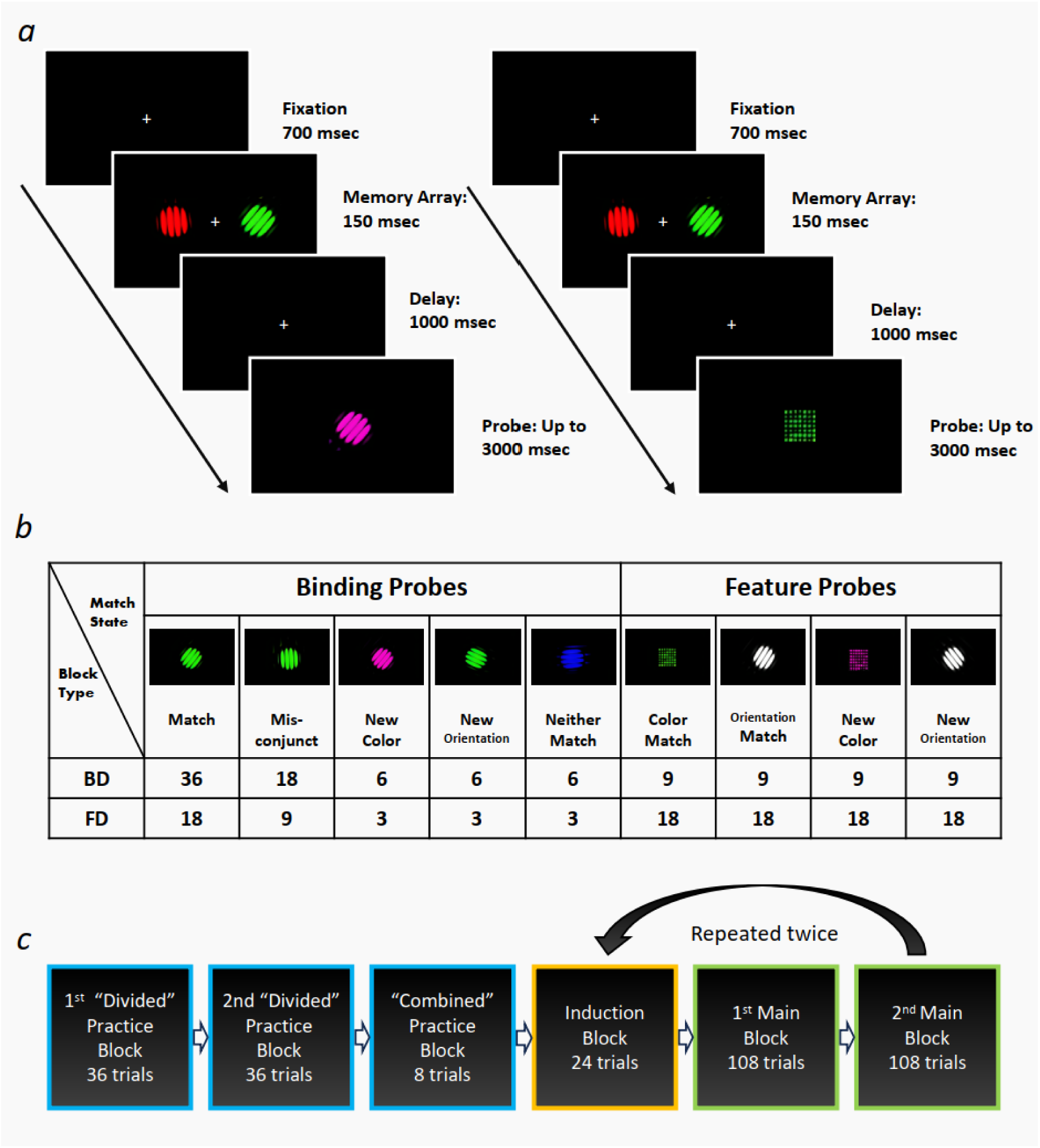
Design of Experiment 1. (a). Left panel-Example of a binding trial with a “Neither match” probe in the main blocks. Right panel-Example of a feature trial with a “color match” probe. (b). Examples of five match states of binding probes and four match states of feature probes labeled in relation to the array shown in panel a. The number represents the number of trials for each match state included in Main block of BD and FD condition respectively. (c). Experimental procedure. Both conditions began with two Divided Practice blocks, one consisting of trials with Binding probes, and the other consisting of trials with Feature probes. This was followed by a Combined practice block, where both types of probes were presented in a random order. Each participant then proceeded to complete two cycles, starting with an Induction block followed by two Main blocks. The Induction blocks consisted of only Binding trials in the BD condition and only Feature probes in the FD condition. In the Main Block, the majority (72 out of 108) of probes in the BD condition were Binding trials, while the majority (72 out of 108) of probes in the FD condition were Feature trials.

### Binding probes

Binding probes can be classified into one of five Match States: Matched, Mis-Conjunct, New-Color, New-Orientation, and Neither-Match (Figure 1b). A Matched probe shared both color and orientation with one of the items in the memory array. A Mis-Conjunct had the color of one of the items in the memory array (“old” color), with the orientation of the other item (“old” orientation). In a New-Color probe, the orientation of one of the items in the memory array was paired with a “New” color that was not present in the array, and vice versa for the New-Orientation probe. Finally, in the Neither-Match probe, both the color and orientation of the probe were not present in the memory array.

### Feature probes

Feature probes contained only one of the two relevant dimensions: color or orientation (Figure 1b). A color probe consisted of an array of small, filled circles within a square 300 pixels in length in either old (Color-Match probe) or new (New-Color probe) colors, presented at the center of the screen. An orientation probe consisted of a colorless gray Gabor grating presented with an orientation either present in the memory array (Orientation-Match probe) or not (New-Orientation probe).

Thus, memory arrays were similar in binding and feature trials, and the appearance of the probe informed the subjects whether they needed to perform color, orientation or color-orientation conjunction recognition.

### Procedure

There were two conditions: Binding dominant (BD) and Feature dominant (FD), which were administered to the same subject on separate days (within-subject design). In the BD condition, the majority of trials presented Binding probes, and in the FD condition, the majority of trials presented Feature probes. The order of conditions was counterbalanced between subjects, and the order of trials was randomized within each subject.

Each condition consisted of four block types: two Divided Practice Blocks, one Combined Practice Block, one Induction Block and two Main Blocks. The combination of one Induction Block with two Main Blocks was repeated twice (Figure 1c).

### Divided Practice Blocks

The experiment began with two practice blocks with 36 trials each. One block included only trials with Binding probes, while the other consisted of only trials with Feature probes. The order of these two practice blocks was counterbalanced across subjects to minimize potential order effects. Each practice trial began with a 100-millisecond fixation cross presented at the center of the screen. Next, the memory array appeared for 400 milliseconds, followed by a 1-second blank screen. After the delay period, a probe item was presented, and participants had to press the key “J” to indicate that the probe item was a “Match” (i.e., present in the memory array) or the key “F” to indicate that the probe item was a “non-match.” The probe disappeared when the response was made, or 3 seconds elapsed. Feedback was then given for 300 milliseconds, indicating whether the response was correct, incorrect, or too late (>3 sec).

### Combined Practice Block

Following the Divided practice blocks, participants proceeded to a Combined Practice block. The trial procedure in this block was the same as that in the Divided Practice Blocks. However, this block presented both types of probes: Four Binding probe trials and four Feature probe trials, in random order. By exposing participants to a mix of probe types, the Combined practice block familiarized subjects with switching between different types of probes.

### Induction Block

After the practice phase, participants engaged in an Induction block with 24 trials. In the FD condition, the Induction block consisted of only feature probes with six trials for each probe type (Color - match, Orientation-Match, New-Color, New-Omrientation). In the BD session, the induction block consisted of only binding probes (12 Matched probe; 6 Mis-Conjunct probes; and 2 each of New-Color, New-Orientation, and Neither match probes). The aim of this block was to induce a specific maintenance format before the main testing blocks began.

### Main Block

Upon completion of the Induction block, participants proceeded to two main testing blocks. The trials in these blocks had the same structure as those in the practice blocks except that the feedback was replaced by a 200-msec fixation cross and that the maximum time for the subjects to respond was reduced to 2 seconds (Figure 1a, b). Each main block consisted of 108 trials. In the FD condition, there were 72 feature trials and 36 binding trials (see Figure 1c for the number of trials for each match state). In the BD condition, there were 72 binding trials and 36 feature trials (Figure 1c).

### Analysis

Statistical analysis was conducted with JASP (version 0.17.1; https://jasp-stats.org/), and figures were made with the python graphic library Seaborn (https://www.python-graph-gallery.com/seaborn/). The proportion of correct responses and reaction times for correct trials were used as the dependent variables. The average response accuracy was calculated for each of four combinations made of two Probe Types (Binding probe vs Feature probe) and two conditions (BD and FD). Subjects with an accuracy below chance (50%) for two of these four combinations were excluded from further analysis under the premise that these subjects did not understand or pay attention to the task.

Accuracy: The Shapiro‒Wilk test showed that the accuracy data were not normally distributed; therefore, we used a data-driven permutation test to determine the significance of the accuracy data (Anderson and Ter Braak, 2003; Suckling and Bullmore, 2004). We performed a Condition (BD, FD) × Probe Type (Binding, Feature) 2×2 repeated-measures ANOVA with the permutation method. Each main effect and interaction were tested separately. For example, to construct the null distribution of the main effect of Condition, the labels BD or FD were randomly permuted within each type and each subject 10000 times. Building up the null distribution of the interaction effect, on the other hand, involved random permutation of the label across both factors within each subject (Anderson & Ter Braak, 2003). F values from the true data larger than 95% of the F values in the permuted distribution indicated a significant effect that is unlikely due to chance. When an interaction was significant, simple effects for each probe type were tested with the Wilcoxon signed-rank test and Bayesian Wilcoxon signed-rank test with 5000 permutations.

Reaction time: The paradigm emphasized accuracy over RT by giving the subject a relatively long time to respond. Therefore, the RT analysis focused on determining whether any interaction found for accuracy was due to the speed-accuracy trade-off. All RTs in this study were normally distributed (Shapiro and Wilk, 1965); hence, we analyzed RTs with a Condition (BD, FD) × Probe Type (Binding, Feature) 2×2 repeated-measures parametric ANOVA. If interactions were found, simple effects were tested with paired-sample t tests and Bayesian paired-sample t tests. In this case, the prior distribution for Bayes analysis was assigned a Cauchy prior distribution (Ly, Verhagen, & Wagenmakers, 2016; Rouder, Speckman, Sun, Morey, & Iverson, 2009) with r= 1/√2(van Doorn et al., 2019).

Our first hypothesis was that performance on different probe type (Binding probe vs Feature probe) would be influenced by the task relevance of conjunction, which would be manipulated through the dominance of probe type within a Condition (BF vs FD block). Thus, we predicted an interaction between Condition (BD, FD) and Probe Type (Binding probe, Feature probe).

Our second hypothesis was that differences in the performance of the binding probe between the BD condition and FD condition would be particularly apparent for Mis-conjunct probes but not for the other Match States. This is because remembering a list of all features in the array could lead to correct performance for all match states except the Mis-conjunct probe if subject react “Non-matched” only when a new feature was introduced (the reader is invited to check this on Figure 1b). If task (induced by different probabilities of probe type) affects the mode of retention beyond any practice/surprise effects, we predicted that the condition effect will be mostly found in Mis-conjunct probe.

To test this prediction, we compared the performance in the BD and FD conditions for each match state of binding trials with paired-sample t tests or Wilcoxon signed-rank tests when the normality assumption was not satisfied. Since we predicted no differences in some of the comparisons, we coupled this analysis with Bayesian paired-sample t tests with a Cauchy distribution with r= 1/√2 as the prior distribution (Ly et al., 2016; Rouder et al., 2009; van Doorn et al., 2019) or the Bayesian Wilcoxon signed-rank test with 5000 permutations when the normality assumption was not satisfied. To ensure that the effect of Condition on the performance of the Mis-conjunct probes was greater than that for the other match states, we compared the differences between condition (BD vs FD) in the performances of the probes on the mis-conjunct probes against all the other four match states (Match; New-Color; New-orientation; Neither-Martch probe).

The datasets and analysis for this study are accessible on OSF (Open Science Framework) at the following link: https://osf.io/w2s84/

### Results and Discussion

A group of subjects performed two conditions of a delayed recognition (yes/no) task in which they decided whether a probe item belonged to a previously presented array of items (Figure 1a). In these two conditions, the proportion of Binding probes and Feature probes was manipulated. As only the Binding probe, required the memory of conjunction between features, we surmised that the conjunction between features is more task-relevant in the condition where the Binding probe was frequent (binding dominant, BD) than when the Feature probe was frequent (feature dominant, FD)

### The effect of task relevance for feature and binding probes

A Condition (BD, FD) × Probe Type (Binding, Feature) 2×2 repeated-measures permutation ANOVA conducted on accuracy data revealed a significant interaction (p=0.001 by permutation, see methods section). A follow-up Wilcoxon test revealed better performance for the BD condition than for the FD condition for Binding probes, *W=205, z=2.55, p=0.011, BF₁₀=8.32,* with no differences between conditions for Feature probes, *W=117.5, z=-0.905, p=0.365, BF₁₀=0.33. T*he permutation ANOVA also suggested that the accuracy of the feature probes was greater than that of the binding probes (main effect of probe type p <0.001), without a main effect of condition type (p=0.23).

For RTs, a Condition Type (BD,FD) × Probe Type (Binding vs Feature) 2 × 2 repeated-measures ANOVA revealed no significant main effect for Probe type, *F(1,23)= 0.11, p =0.74,* and a main effect of condition, with RT being shorter in the FD condition than in the BD condition, *F(1,23)= 37.32, η2 = 0.06, p < .001*.

The effect of condition was significant for both Feature probes, *t(23) = -7.16, p < .001, Cohen’d = -1.46, B10 = 59114.9*, and Binding probes, *t(23) = 4.33, p < .001, Cohen’d = 0.88, B10 = 119.74; however*, the effect of condition was greater for feature probes, as indicated by a significant interaction (*Figure 3b*; *F(1,23) =51.02, η2 = 0.06, p < .001)*.

To conclude, these results supported our first hypothesis that performance on different probe types (Binding vs Feature probes) was influenced by different levels of task relevance of conjunction, which was manipulated through the dominance of probe type within a condition (BF vs FD condition).

Specifically, responding to binding probes became more accurate when these probes were more abundant, and RTS for feature probes was faster when these probes were abundant, without affecting accuracy.

### The effect of the task on each type of binding probe

We hypothesized that maintenance of conjunctions will be critical predominantly for correctly recognizing the Mis-conjunct probe, while all other probes could be judged correctly by maintaining separate features, and identifying each probe with new features as ‘non-match’ (e.g., a color absent from the array), and as ‘match’ when no new feature is detected. However, as there are no new features of the Mis-Conjunct probe, this strategy could result in mistakenly identifying a Mis-conjunct probe as a “match”. Therefore, among all the match states in the binding probe, we predicted that the Mis-Conjunct probe would be the most sensitive to the induction task. Indeed, the planned contrast showed that the effect of condition (BD condition-FD condition) on accuracy was greater for the Mis-Conjunct probe than for the other probes combined, *t(92) = 2.63, p = .01* (Figure 2c). That is, the Mis-conjunct probes benefited from the binding probes being more frequent than other match states. The Wilcoxon signed-rank test was subsequently applied to compare each match state in terms of their condition effects. Only the Mis-conjunct probes elicited a significant effect on Condition (Table 1a). These results also provided evidence that the better performance of binding probe in BD than FD condition we found was unlikely to be due to practice-related improvement in response to the binding probe in BD condition or surprise-related impairment in response to the binding probe in FD condition, as all binding probes were more frequent in the BD condition than in the FD condition.

**Table 1a.**
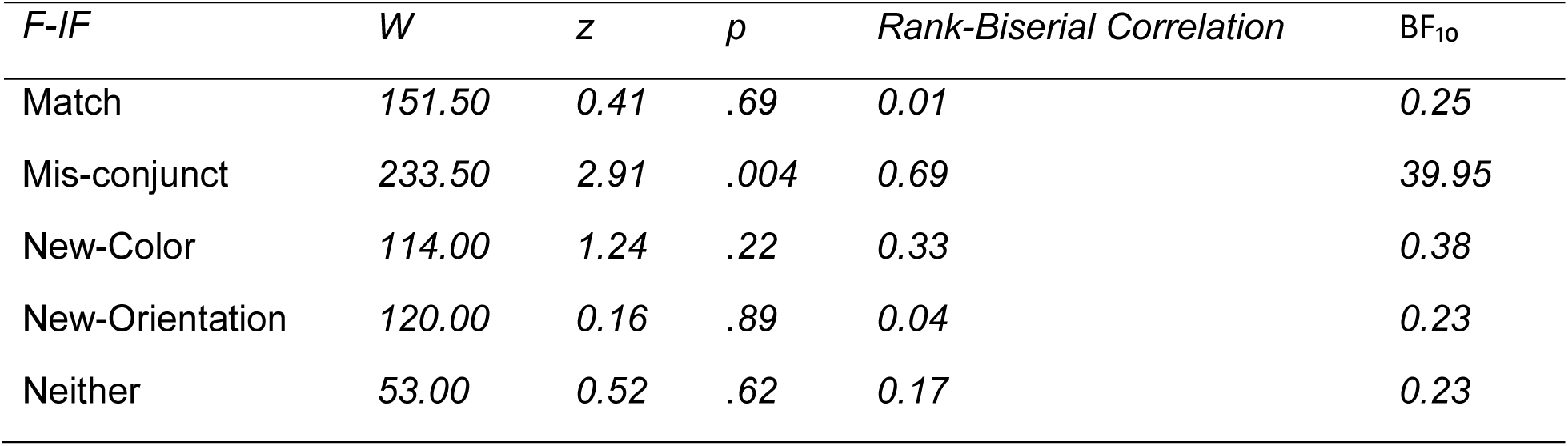
Effect of Task on Each Match State Binding Probe on the ACC.

| *F-IF* | *W* | *z* | *p* | *Rank-Biserial Correlation* | *BF<sub>10</sub>* |
| --- | --- | --- | --- | --- | --- |
| Match | 151.50 | 0.41 | .69 | 0.01 | 0.25 |
| Mis-conjunct | 233.50 | 2.91 | .004 | 0.69 | 39.95 |
| New-Color | 114.00 | 1.24 | .22 | 0.33 | 0.38 |
| New-Orientation | 120.00 | 0.16 | .89 | 0.04 | 0.23 |
| Neither | 53.00 | 0.52 | .62 | 0.17 | 0.23 |

**Figure 2.**
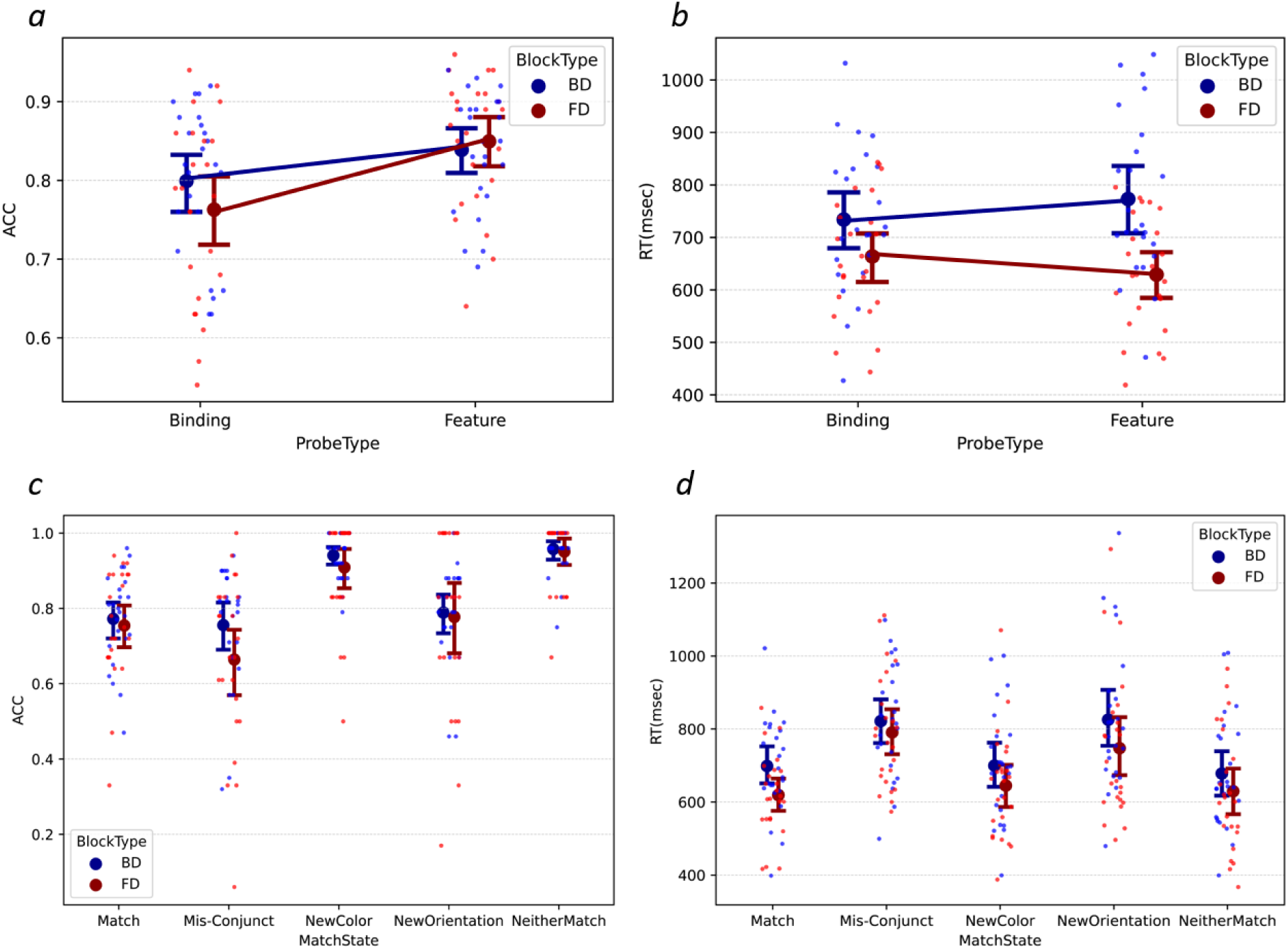
Results of experiment 1. *Note (*a)-(b). Average response accuracy (a) and reaction times (b) for binding and feature probes from the BD and FD conditions. Here, and in other panels, small dots represent individual subject means; large dots and error bars depict across-subjects means and 95% confidence intervals. (c)-(d): Response accuracy (c) and reaction time (d) under BD and FD conditions in each match state for the binding probe.

**Figure 3.**
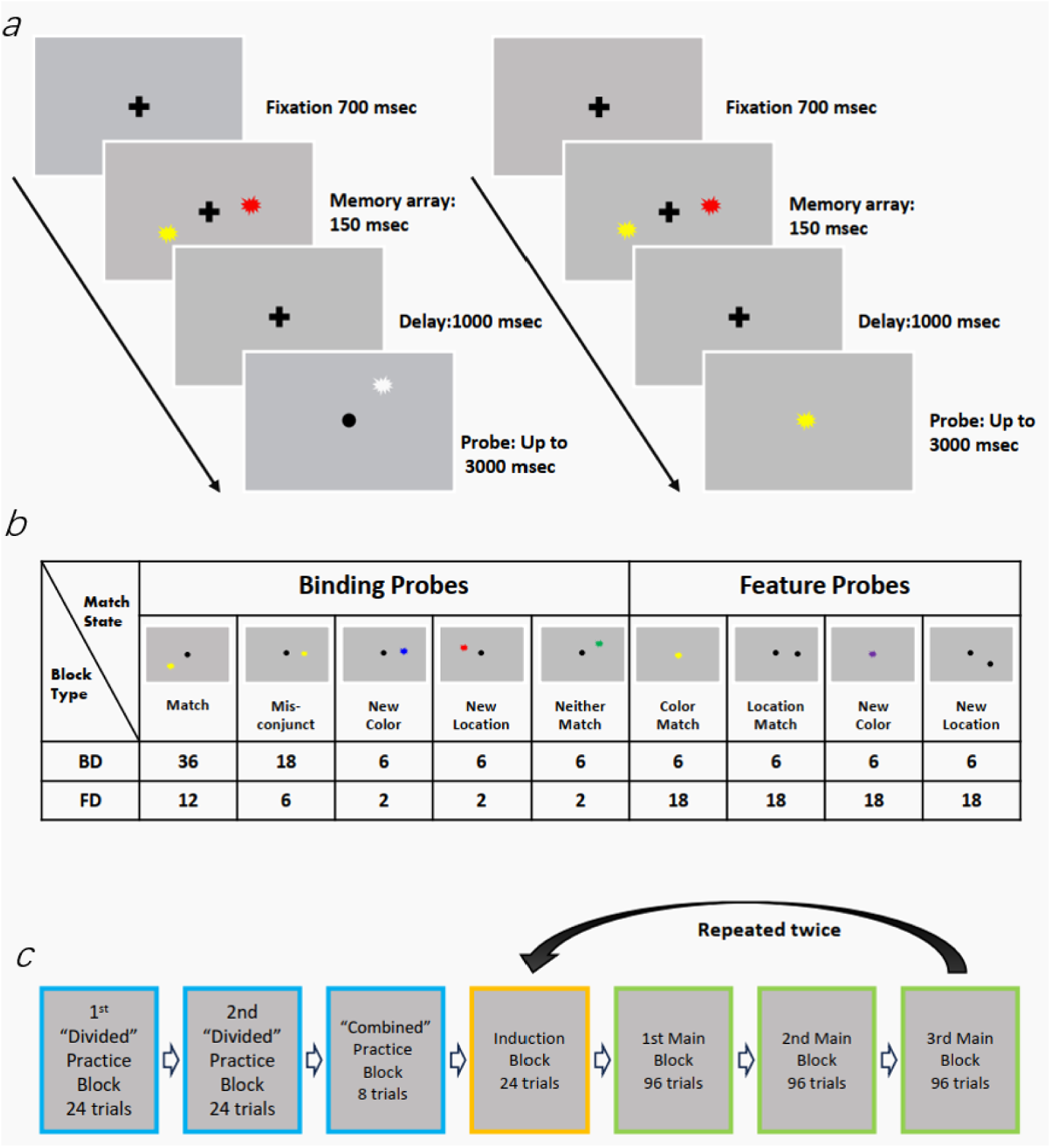
Design of Experiment 1. *Note.* (a): Left panel-An example of a binding trial showing a Neither-Match binding probe; right panel-An example of a feature trial showing a Color-Match probe. (b): Examples of five match states of the binding probe and four match states of the feature probe labeled in relation to the array shown in panel a. (c): Experimental procedure. Both conditions began with two Divided practice blocks, one consisting of trials with Binding probes, and the other consisting of trials with Feature probes. This was followed by a Combined practice block, where both types of probes were presented in a random order. Each participant then completed two cycles, starting with an Induction block followed by three main blocks. The Induction blocks consisted of Binding trials in the BD condition and Feature probes in the FD condition. The majority (72 out of 96) of probes in the Binding conditions were Binding trials, while the majority (72 out of 96) of probes in the Feature conditions were Feature trials.

As reported above, RTs were overall faster in the FD condition than in the BD condition. If the higher accuracy in the BD condition compared to the FD condition for the Mis-Conjunct probe was due to a time-accuracy trade-off, we would expect to see that the Mis-Conjunct probe would perform more slowly in the BD condition, increasing the overarching RT difference between the BD and the FD conditions compared to other match states. The planned contrasts did not reveal such a pattern; rather, the opposite was found—the difference between the BD and FD in RTs was marginally smaller for the Mis-Conjunct probe than for the other match states, *t(92) = -1.745, p = .084 (Figure 2d).* Table 1b shows the Wilcoxon signed-rank tests for each match state. The BD condition responded more slowly than did the FD condition for binding probes in all match states but not for the Mis-Conjunct. These results suggest that the greater accuracy of the Mis-conjunct probe in the BD condition than in the FD condition was unlikely due to the time-accuracy trade-off.

**Table 1b.** Effect of Task on Each Match State Binding Probe on the RTs.

| *FI-F* | *t* | *df* | *p* | *Cohen's d* | *BF<sub>10</sub>* |
| --- | --- | --- | --- | --- | --- |
| Match | 4.559 | 23 | < .001 | 0.931 | 200.457 |
| Mis-conjunct | 1.475 | 23 | 0.154 | 0.301 | 0.557 |
| New-Color | 3.258 | 23 | 0.003 | 0.665 | 11.925 |
| New-Orientation | 2.859 | 23 | .009 | 0.584 | 5.356 |
| Neither | 2.073 | 23 | 0.050 | 0.423 | 1.310 |

To conclude, we found that memory was better for Binding probe in the BD condition than in the FD condition, supporting our hypothesis that maintaining color and orientation as separate features, or their conjunction depends on the task. The Mis-conjunct probe showed the effect of condition more than other binding probes in the same condition indicating that such task effect is not due to general practice. Additionally, we found that the trials in the BD condition were performed slower than those in the FD condition, suggesting that maintaining separate features might be easier than maintaining conjunctions.

### Experiment 2

Experiment 1 revealed a flexibility of working memory maintenance for the conjunctions of color and orientation, two features that are not location specific. Experiment 2 aimed to examine the flexibility of maintaining conjunctions between color and spatial location. Similar to Experiment 1, this study manipulated the relevance of conjunctions and separate features and predicted an interaction between Condition (BD vs FD) and Probe type (Feature probe vs Binding probe). We hypothesized that the effect of task relevance would be observed mostly on Mis-Conjunct probes, which particularly require the maintenance of conjunctions.

## Materials and methods

### Participants

Power analysis based on Experiment 1 suggested that fifteen participants were required to reach 80% of the power with an alpha of 0.05 for the critical contrast between Mis-conjunct probes in the BD and FD conditions. Seventeen students, recruited through the Hebrew University online study recruitment system, participated in the study (age range 21-35 years, mean± SD =25.02 ± 2.90 years, 77% females). Subjects with any two of four combinations (Binding Probe in BD condition, Binding Probe in FD condition, Feature probe in BD condition, Feature probe in FD condition) below a response accuracy of 0.5 were excluded, resulting in the exclusion of two participants. The study was approved by the ethics committee of the Social Sciences Faculty at the Hebrew University of Jerusalem, Israel. The subjects downloaded the computer program made with E-Prime 3 online (https://pstnet.com/eprime-go/) to their own computers. Each participant conducted two conditions of the experiment on different dates within a week. They received either two course credits or a monetary compensation of 60 NISs (∼$15) for their participation.

### Stimuli

The stimuli consisted of spiky colored blobs, presented in memory arrays of two items (Figure 3a). The color of each item was randomly selected out of six highly distinguished colors, including red (RGB: 255,0, 0), green (0, 255, 0), yellow (255,255, 0), blue (0, 0, 255), violet (255,255,0) and white (0,0,0). The same color never appeared twice in an array. The exact visual angle and size of the stimulus varied as participants performed the tests on their own screens. Each spiky colored blob was limited within an invisible square of 55 x 55 pixels. The two selected locations were randomly selected out of six potential locations evenly distributed on an invisible circle with a diameter of 140 pixels centered on the fixation cross. The stimuli were created with Psychotoolbox-3, implemented in MATLAB 2018. Each probe contained only one item of the same size as the items presented on the memory array. There were two types of probes: Binding probes and Feature probes (Figure 3b):

### Binding probes

The probes consisted of a spiky colored blobs presented at one of the potential locations used in the memory arrays. To answer correctly, the subjects had to maintain the conjunction information and answer whether the probe was the same as an item from the array in both color and location. Binding probes (Figure 3b) included five match states (Match, Mis-Conjunct, New-color, New-location, and Neither-Match) according to their relation to the previous memory array with the same principle detailed in Experiment 1.

### Feature probes

Feature probes (Figure 3b) consisted of either one black item at one of the possible locations (a location probe) or a colored item presented at the center of the screen (a color probe). Subjects were required to indicate whether location probes appeared in the same location as one of the items in the memory array (Location-match) or not (New-Location), whereas for color probes, subjects were required to indicate whether the color of the item appeared in the memory array (Color-match) or not (New-color), regardless of its location. Thus, retention of separate features in working memory would allow a more effective response for feature probes than retention of conjunctions (which would entail a need to disconnect a stimulus from its bound location).

### Procedure

As in experiment 1, there were two conditions: Binding dominant (BD) and Feature dominant (FD), administered to the same subject on separate days (within-subject design). In the BD condition, the majority of trials presented Binding probes, and in the FD condition, the majority of trials presented Feature probes. The order of conditions was counterbalanced between subjects, and the order of trials was randomized within each subject. Each condition consisted of four block types: two Divided Practice Blocks, one Combined Practice Block, one Induction Block and two Main Blocks. The combination of one Induction Block with two Main Blocks was repeated twice (Figure 3c).

### Divided Practice Blocks

The experiment began with two practice blocks of 24 trials. One block included only trials with Binding probes, while the other block consisted of only trials with Feature probes. The order of these two practice blocks was randomized across participants. Each practice trial began with a 700-millisecond fixation cross presented at the center of the screen. Next, the memory array appeared for 150 milliseconds, followed by a 1-second blank screen. After the delay period, a probe item was presented, and participants had to press the key “J” to indicate that the probe item was “Match” (i.e., present in the memory array) or the key “F” to indicate that the probe item was “Non-match.” The probe disappeared when the response was made or 3 seconds elapsed. Feedback was then given for 300 milliseconds, indicating whether the response was correct, incorrect, or too late.

### Combined Practice Block

Following the Divided Practice blocks, participants proceeded to a Combined Practice block. The trial procedure in this block was the same as that in the Divided Practice Blocks. However, this block presented both types of probes—four binding probe trials and four feature probe trials—in random order. By exposing participants to a mix of probe types, the combined practice block familiarized subjects with switching between different types of probes.

### Induction Block

After the practice phase, participants engaged in an Induction block of 24 trials. The aim of the Induction block was to induce a specific maintenance format before the testing blocks began. Trials in induction blocks had the same structure as those in the Divided Practice block. In the FD condition, the induction block consisted of only feature trials with six trials for each probe type (Location-Match probe, New-Location probe, Color-Match probe, New-Color probe). In the BD session, the induction blocks consisted of only binding probes (12 Matched probe; 6 Mis-Conjunct probes; 2 for New-Color, New-Location, and Neither match).

### Main Block

Upon completion of the Induction blocks, the participants proceeded to three main blocks. The trials in the main blocks had the same structure as those in the practice blocks, except that the feedback was replaced by a 100-msec fixation cross, and the maximum time for the subjects to respond was reduced to 2 seconds (Figure 3a). Each testing block consisted of 96 trials. In the FD condition, there were 72 feature trials and 24 binding trials, whereas in the BD condition, there were 72 binding trials and 24 feature trials (see Figure 3b for the number of trials for each match state).

### Analysis

The analysis was the same as that in Experiment 1. The datasets and analysis for this study are accessible on OSF (Open Science Framework) at the following link: https://osf.io/w2s84/

## Results

The subjects performed a task similar to that of Experiment 1 except that the two relevant features were the color and location of the stimuli. The task relevance of conjunctions was manipulated as before by manipulating the proportion of binding probes and feature probes.

### The effect of task relevance on binding and feature probes

Condition (BD, FD) × Probe Type (Binding probe, Feature probe) 2 × 2 repeated-measures permutation ANOVA was conducted on accuracy with 10000 permutations *(Figure 4a)*. Binding probes were performed significantly better in the BD condition than in the FD condition, while Feature probes were performed significantly better in the FD condition than in the BD condition, resulting in a significant interaction, *p <.001* by permutation, with neither the main effect for the Probe Type (*p=.25*) nor for Condition (*p = .34*)

**Figure 4.**
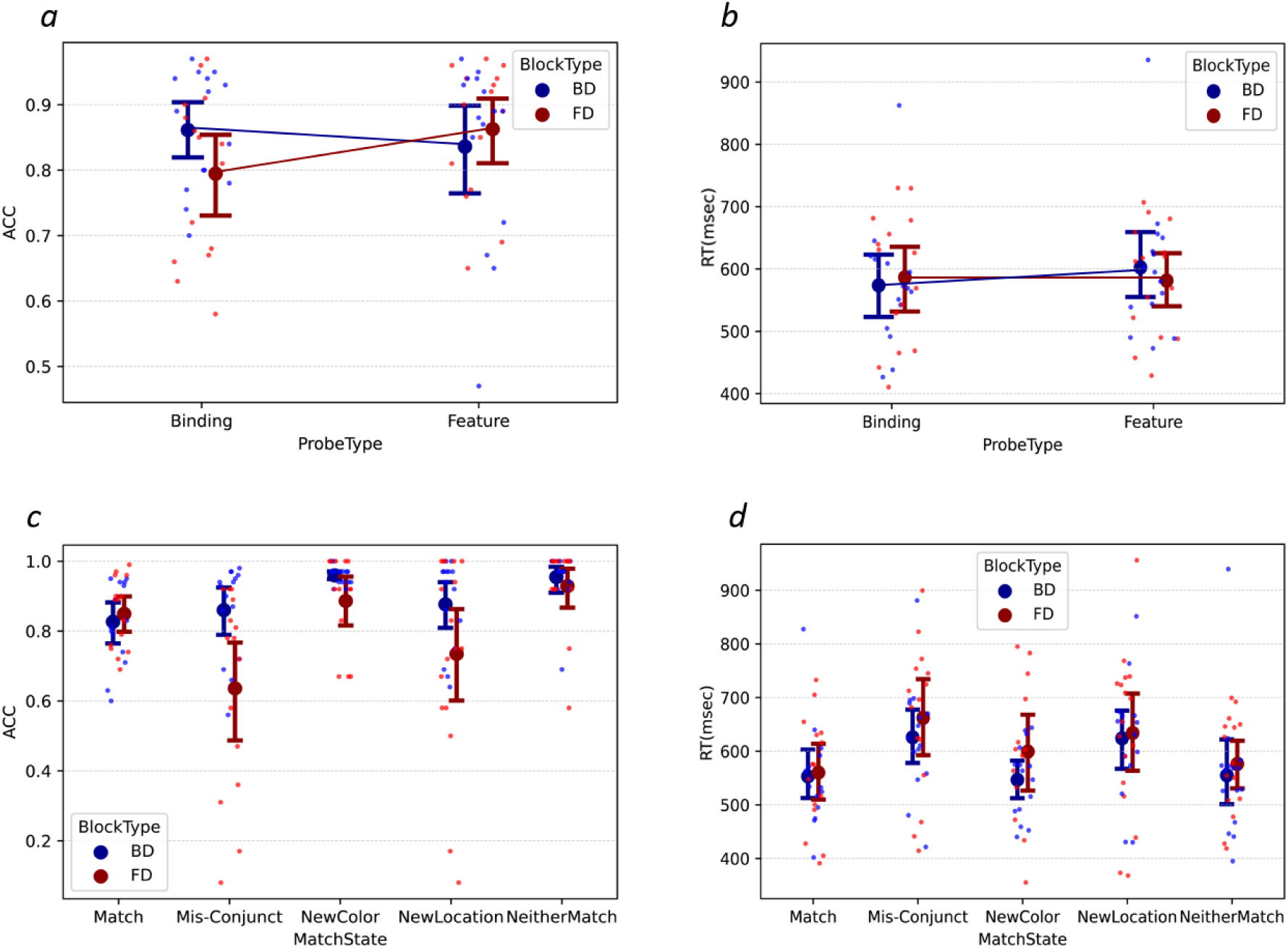
Results of Experiment 2. *Note (***a)-(b)**. Average response accuracy (a) and reaction times (b) for binding and feature probes from BD and FD blocks. (c)-(d): Response accuracy (c) and reaction time (d) in the BD and FD blocks in each match state for Binding Probe. The symbols and notations are the same as those in Figure 2.

A Condition (BD, FD) × Probe Type (Binding, Feature probe) 2 × 2 repeated-Repeated measures ANOVA revealed a somewhat faster response for Binding probes in BD than in FD condition and for Feature probes in FD than in BD condition *(Figure 4b*), resulting in a significant interaction between Condition and Probe type, *F(1,14) = 17.03, p = .001, η2 = 0.03*. In addition, subjects responded faster to Binding probes than feature probes (Main effect of Type), *F(1,14) =5.43, p = .04, η2 = 0.01*, while no significant main effect of condition was found, *F(1,14) =0.02, p = .88*.

Thus, both Feature and Binding probes showed faster and more accurate memory performance when presented in the corresponding task-relevant condition, which supported our hypothesis that task demand could influence the retention formats of objects in working memory.

### The effect of task relevance on each type of binding probe

We next split the Binding probe trials to examine the effect of task relevance for each Match State of probe. As in Experiment 1, our flexible representation hypothesis predicted that larger differences between probes in the BD and FD conditions would be found for the Mis-Conjunct probe trials than for the other match states. Consistently, the planned contrast between the Mis-Conjunct probe and all other match states combined showed that the condition effect of the Mis-Conjunct probe on accuracy was greater than that of the other match states combined*, t(56) = 3.38, p <.001*. The Wilcoxon signed-rank test was further applied to compare the response accuracy for trials in the BD and FD conditions for each match state. As expected, the accuracy of the Mis-Conjunct probe was greater when the binding probes were tested frequently under BD conditions than under FD conditions. Unexpectedly, the same effect was shown for the New-Location probe *(Table 2a)*.

**Table 2a.**
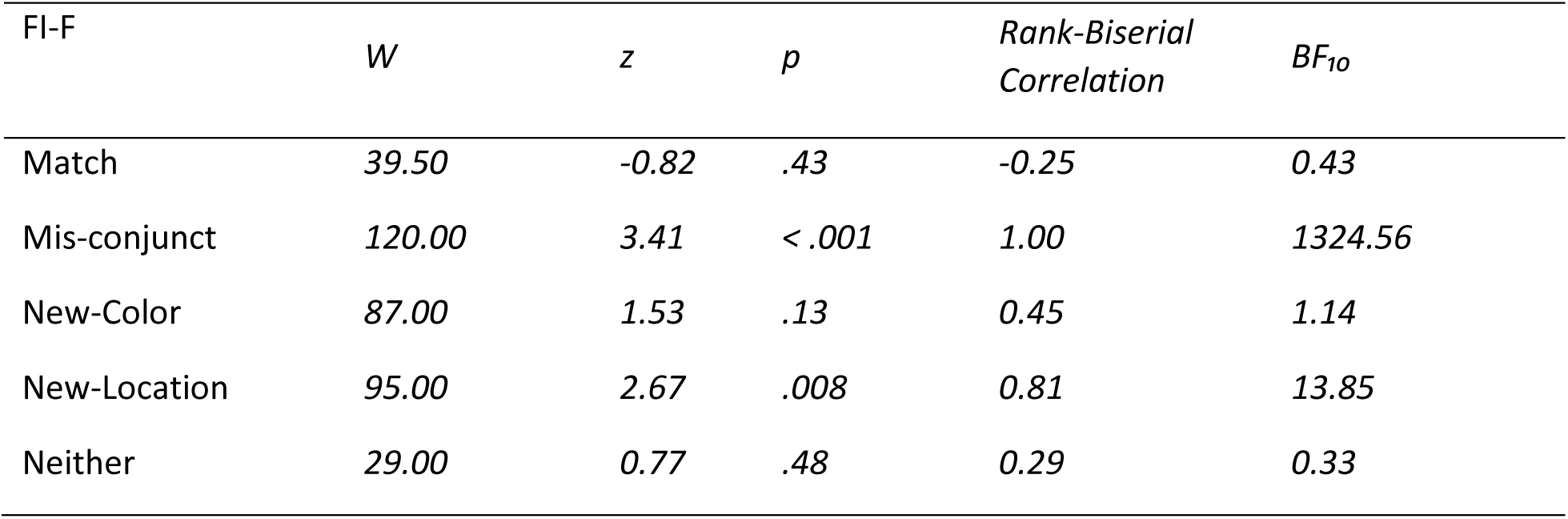
Effect of Task on Each Match State Binding Probe on the ACC.

| FI-F | *W* | *z* | *p* | *Rank-Biserial Correlation* | *BF<sub>10</sub>* |
| --- | --- | --- | --- | --- | --- |
| Match | 39.50 | -0.82 | .43 | -0.25 | 0.43 |
| Mis-conjunct | 120.00 | 3.41 | < .001 | 1.00 | 1324.56 |
| New-Color | 87.00 | 1.53 | .13 | 0.45 | 1.14 |
| New-Location | 95.00 | 2.67 | .008 | 0.81 | 13.85 |
| Neither | 29.00 | 0.77 | .48 | 0.29 | 0.33 |

For all the match states, the responses to Binding probes were nominally faster in the BD condition than in the FD condition, but these differences were not significant, nor was the planned contrast between the differences in the BD and FD conditions for the Mis-Conjunct probes and the other four Match States combined, *t(56) = 1.08, p= .29 (Table 2b)*. This suggests that the difference in response accuracy between Match States was not due to a tradeoff between speed and accuracy.

**Table 2b.** Effect of Task on Each Match State Binding Probe on the RTs.

| FI-F | *t* | *df* | *P* | *BF<sub>10</sub>* |
| --- | --- | --- | --- | --- |
| Match | 0.28 | 14 | .78 | 0.27 |
| Mis-conjunct | 1.12 | 14 | .28 | 0.45 |
| New-Color | 1.00 | 14 | .33 | 0.40 |
| New-Location | 0.26 | 14 | .80 | 0.27 |
| Neither | 0.74 | 14 | .47 | 0.33 |

To summarize, this experiment showed that the response accuracy was lower when a Mis-Conjunct probe was used in the FD condition, where the binding probes were presented infrequently, and that the response accuracy was greater for the BD when the binding probes were frequent; thus, maintaining conjunctions was critical. An unexpected finding was that response accuracy for new location probes was also impacted in the FD condition when conjunction information was less task relevant. In other words, when subjects were not required to report on the binding between color and location, location information tended to be lost. Comparing Experiments 1 and 2, maintaining features seemed to be easier than maintaining conjunctions between color and orientation (Experiment 1), whereas for conjunctions between color and spatial location (Experiment 2), this advantage was not observed, and in fact, the Binding probes elicited faster responses relative to the Feature probes, without compromising accuracy. To confirm this observation statistically, we compared the Feature advantage on ACC (*Frequent Binding probe-Frequent Feature probe*) and RTs (*Frequent Feature probe-Frequent Binding probe*) between Experiment 1 and Experiment 2. As expected, we found a significantly greater advantage for features (color or orientation) in Experiment 1 than for those with color and location (Experiment 2), both in the ACC, *t(37) = 2.42, p=.021, Cohen’d = 0.36,* and in RTs, *t(37) = 4.14 p < .001, Cohen’d = 0.41.* This suggests that in addition to task relevance, the involvement of specific stimulus dimensions may play a role in determining representation formats.

### General Discussion

In this study, we investigated whether the task determines the maintenance format in visual working memory—as separate features or as bound objects. In two studies using a delayed (yes/no) recognition task, we implicitly biased subjects to maintain features or conjunctions by manipulating the proportion of probes that required conjunction information. Our results from the two studies show that both trials that required feature-level information and those requiring conjunction information performed better when they were more frequent in a block of trials. Overall, this study provides novel evidence for the flexibility of maintenance formats in VWM based on task requirements.

Previous studies have demonstrated the flexibility of working memory by showing that task-relevant features could be selectively retained while discarding task-irrelevant features presented simultaneously (Park et al., 2017; Pertzov, Bays, et al., 2013; Ye et al., 2016). These observations emphasized the functional purpose of working memory and challenged the view that conceptualized working memory as a task-independent system (Nobre & Stokes, 2020). Our study investigated a different type of flexibility that received less attention in that respect: whether visual information is retained as separate features (e.g., color, orientation) or as bound objects. We presented probes that required conjunction information and probes that did not, and by manipulating the relative dominance of these two probes within a block, we implicitly created a task set favoring one of the two maintenance strategies. If the retention formats were independent of the task context, performance for both types of probes should have remained the same in the two dominance conditions. In contrast, we found that retention formats were shaped by the task requirement: performance with Binding probes was better when these probes were dominant (in the BD condition) than when Feature probes were dominant (in the FD condition), and vice versa for Feature probes. In accordance with our expectations, this effect manifested especially in trials in which Mis-conjunct probes were presented, a type of trial that critically requires the retention of conjunction information. These findings provide a new perspective for understanding some seemingly contradictory results and call attention to task requirements when investigating maintenance formats in VWM.

We are aware of only one study that directly manipulated the relevance of binding. In their study, Vergauwe and Cowan (2015) compared reaction times (RTs) when the subjects were explicitly instructed to search for colors, shapes, or both. They found that retrieval of both features took no longer than retrieval of a single feature when the memory of bound objects was encouraged. Otherwise, RT increased with the number of features to be retrieved. They concluded that whether participants preserved bound objects or separate features in VWM is dependent on the testing situation. Here, we explored the binding between color and location and between orientation and color. We found that even without explicitly instructing subjects about the preferred mode of maintenance, the format of items in working memory will be flexibly adjusted to optimally perform the task.

Whether conjunction memory is required in the task is a factor relevant to all experiments aiming to investigate the object representation formats. Some paradigms have explicitly addressed this aspect. For example, to study whether a location is automatically bound to a nonspatial feature, studies usually ensure, and explicitly report, that the conjunction of spatial and nonspatial features is task irrelevant (Foster et al., 2017). In other studies, the relevance of conjunction for completing a task has not been reported but could be determined from the paradigm. For example, a study by Fougnie and Alvarez (2011) revealed that errors in memory for one feature did not significantly affect memory for other features. It should be noted that in their design, participants were asked to recall either the color or orientation of an object separately, rendering the conjunction information task irrelevant. Whether their observation would hold had conjunction information been task-relevant is unknown. In some studies, the task-relevant information is subtler and cannot be directly determined from the experimental paradigm without careful attention to the experimental details. For instance, in Wheeler and Treisman’s (2002) delay-match-to-sample task, features of different objects were not repeated within a memory array, and the probe either contained a new feature or matched one of the objects kept in working memory in both features, making it unnecessary to retain conjunctions (as explained above, this task can be solved simply by deciding whether the probe presents a new feature or not). In contrast, in Luck and Vogel’s study (1997; extended in Vogel, Woodman, & Luck, 2001), maintaining conjunctions was necessary to correctly respond, as the memory array consisted of items with repeated features (e.g., two objects in an array could be red), and Mis-conjunct probes were occasionally presented (where the probe was a mismatch even though no new feature was presented). Consistent with this design difference, Wheeler and Treisman (2002) concluded that features were maintained separately, whereas Luck and Vogel concluded that features were conjoined in VWM. More investigations are required to test whether such subtle differences lead to different conclusions about retention formats in VWM. Nevertheless, the flexibility of representation formats found in the current study mandates that studies investigating retention formats will explicitly consider the task relevance of conjunctions in their design and investigate the effect of the task along with other variables of interest whenever possible.

Our study additionally suggested that task relevance alone cannot completely determine the representation formats of objects in VWM. Rather, the degree to which an object was kept in one format or the other varied depending on other factors. An exclusive task effect would have been reflected by better performance on Binding probes than Feature probes in the BD condition and better performance for Feature probes than Binding probes in the FD condition. However, this was not what we found. For color and orientation features, the performance of feature probes outperformed that of binding probes (Experiment 1), regardless of whether conjunctions were task relevant. This finding is consistent with studies showing greater cognitive resources required for remembering the conjunction of two features not involving spatial location, including greater involvement of cortical regions (Parra, Della Sala, Logie, & Morcom, 2014) and increased impairment of conjunction memory due to a secondary task (He et al., 2020; Shen, Huang, & Gao, 2015). In contrast, when location (spatial) and color (nonspatial) features were involved (Experiment 2), we observed a different pattern, whereby subjects were faster for Binding probes than for Feature probes regardless of the task relevance of conjunctions. This finding is consistent with studies showing that location can be automatically encoded (Elsley & Parmentier, 2015; Foster et al., 2017; Olson & Marshuetz, 2005). Combining the results from both studies, the results confirm our hypothesis regarding the effect of the task relevance of conjunctions in determining the storage format in working memory and suggest a role for which features are to be encoded.

Specifically, participants had difficulty identifying new locations when the conjunctions were de-emphasized, which was not the case for new colors. That is, memory of color was not dependent on memory for its location, but the memory of location required it to be yoked with color. This observation might be surprising, as the spatial dimension has long been considered a primary dimension that allows us to bind other features and to access our internal representations of objects, both visual (Kondo & Saiki, 2012; Rajsic & Wilson, 2012; Treisman & Zhang, 2006; Wheeler & Treisman, 2002) and auditory (Deouell, Bentin, & Soroker, 2000).

One of the causes of interdependence between location memory and conjunction memory observed in the current study might be difficulty in maintaining an empty location. There is some evidence showing that an empty location (i.e., one without other associated features) is especially difficult to remember. For example, when an object suddenly appeared in a previously empty location, infants at least 8 months old were no surprise (Wynn & Chiang, 1998). In contrast, infants as young as 3 months old showed a surprise response when an item disappeared magically (Baillargeon & Devos, 2016). The above two phenomena could be explained by the inability to encode and maintain an empty location. In this case, no representation is formed in visual working memory before the object appears, and no conflict (or prediction error) occurs when it does. On the other hand, when an object is present at some locations, the memory system keeps the object bound to that location, and the disappearance of the object conflicts with the previous representation in working memory. Difficulties in maintaining empty locations were also found in adults (Csink, Gliga, & Mareschal, 2022).

Our study has several limitations that should be addressed in future research. Since we only applied a single delay duration in our study, we could not investigate the temporal dynamics of the process. Specifically, research is necessary to discern whether the task manipulated the initial encoding into working memory or later retention processes. Second, the formats of object representations in working memory were estimated indirectly, based on the assumption that the number of representation units (features or objects) correspondingly consumes several units of working memory resources. This assumption, however, was challenged by models conceptualizing working memory capacity as a continuous resource rather than as discrete units (Ma, Husain, & Bays, 2014). To accumulate convergent evidence and generalize our results, the flexibility of the formats should be tested with a variety of paradigms without assuming the slot model of working memory (Fougnie et al., 2010).

In summary, our research emphasizes the role of experimental design, even implicit design, in shaping the formats of objects in working memory. It offers new evidence that the way an object is represented in working memory, whether as a distinct feature or as an integrated object, varies depending on the demands of the specific task at hand. These findings highlight the adaptability of representation formats beyond the task relevance of any specific feature.

## Acknowledgments

We thank Yoni Pertzov for inspiration on the design and discussion of the present study. We thank Areej Mousa and Neta Licht for helping with the data collection.

